# LiBis: An ultrasensitive alignment method for low-input bisulfite sequencing

**DOI:** 10.1101/2020.05.14.096461

**Authors:** Yue Yin, Jia Li, Jin Li, Minjung Lee, Sibo Zhao, Linlang Guo, Jianfang Li, Mutian Zhang, Yun Huang, Xiao-Nan Li, Deqiang Sun

## Abstract

The cell-free DNA (cfDNA) methylation profile in liquid biopsies has been utilized to diagnose early-stage disease and estimate therapy response. However, in typical clinical settings, only very small amounts of cfDNA can be purified. Whole-genome bisulfite sequencing (WGBS) is the gold standard to measure DNA methylation; however, WGBS using small amounts of fragmented DNA introduces a critical challenge for data analysis, namely a low mapping ratio. This, in turn, generates low sequencing depth and low coverage for CpG sites genome wide. The lack of informative CpGs has become a bottleneck for the clinical application of cfDNA-based WGBS assays. Hence, we developed LiBis (Low-input Bisulfite Sequencing), a novel method for low-input WGBS data alignment. By dynamically clipping initially unmapped reads and remapping clipped fragments, we judiciously rescued those reads and uniquely aligned them to the genome. By substantially increasing the mapping ratio by up to 88%, LiBis dramatically improved the number of informative CpGs and the precision in quantifying the methylation status of individual CpG sites. The high sensitivity and cost effectiveness afforded by LiBis for low-input samples will allow the discovery of genetic and epigenetic features suitable for downstream analysis and biomarker identification using liquid biopsy.

## Introduction

DNA methylation abnormalities contribute to tumorigenesis and tumor prognosis (1). Tumors are characterized with global hypomethylation and focal CpG island hypermethylation (2). To develop a simple, economical assay for cancer early diagnosis or monitoring response to therapy, liquid biopsies have rapidly emerged as an alternative to tumor biopsy due to its minimal invasiveness and ease of repeat sampling (3). Many studies have reported that tumor DNA methylation status can be accurately detected in circulating cell-free DNA (cfDNA) from blood or other body fluids (4-6).

Compared to the limited number of CpGs detected using microarray technology, whole genome bisulfite sequencing (WGBS) can detect all the CpG sites in the genome, which dramatically increases the power for biomarker discovery. However, the extremely low amount of cfDNA in liquid biopsies poses a challenge for whole genome bisulfite sequencing using cfDNA. The amount of cfDNA collected from the plasma of healthy individuals or patients usually ranges from a few nanograms to a few dozen nanograms, and for late-stage cancer patients the amount is usually less than several hundred nanograms, which is still low for WGBS library preparation (7,8). To prepare WGBS libraries using low amounts of DNA, several methods were developed. Post bisulfite sequencing methods such as post-bisulfite adaptor tagging (PBAT) and single cell WGBS methods apply adaptor tagging after bisulfite treatment to reduce the loss of tagged DNA fragments during the library preparation (9). Random primers are designed to capture all single strand DNA generated by bisulfite treatment which increases the sensitivity of library preparation for low input WGBS. However, it also introduces long synthetic sequences by dNTP during the random priming process (10,11). The sequencing reads with variable lengths of synthetic sequence contamination limit the effectiveness of current fixed-length trimming methods (11). Moreover, random primers may also introduce unmappable chimeric reads by concatenating two genomic DNA fragments (12). These issues of WGBS library preparation from low amounts of DNA result in low mapping ratios, which increase the cost of liquid biopsy and restrict its wide application in clinical settings.

Several mapping programs such as BSMAP, Bismark, BS-Seeker, and scBS-map have been developed to address the above issues (13-15). Although BS-Seeker utilizes soft-clipping during the mapping procedure and scBS-map adopts a local alignment strategy to improve the mapping ratios, the mapping ratios of low input WGBS samples are still far lower than traditional WGBS samples (16). To get enough data coverage for all CpGs from low-input bisulfite sequencing data, it is imperative to develop a mapping procedure to recover as much data as possible from the “unmapped” reads due to the library preparation protocol drawbacks. Here, we developed LiBis to further improve the mapping ratio of low-input bisulfite sequencing data. LiBis applies a dynamic clipping strategy to rescue the discarded information from each unmapped read in end-to-end mapping.

In our simulation study, LiBis achieved the highest mapping ratio improvement and the shortest CPU time among published methods. LiBis also improved the number of detected CpGs and the methylation ratio accuracy in a time-efficient manner. By applying LiBis, we achieved better cost efficiency using both a public dataset and in-house datasets. The number of informative CpGs increased significantly after using LiBis compared with using a traditional trimming protocol. The precision of bisulfite sequencing was also improved by LiBis for all samples. Furthermore, LiBis was able to identify virus insertion sites in a cervical cancer WGBS dataset, which indicates that bisulfite sequencing data can be used to reveal both genetic and epigenetic changes. LiBis supports a one-command solution for quality control, trimming, mapping, and methylation calling in a reasonable computing time, making it an effective and comprehensive solution to support large-scale single-cell or cfDNA bisulfite sequencing applications.

## Materials and Methods

### Patient sample collection

Signed informed consent was obtained from all patients or their legal guardians prior to sample acquisition in accordance with an institutional review board-approved protocol. CSF samples were obtained from two brain tumor patients at Texas Children’s Hospital at the time of clinically indicated lumbar puncture. CSF was processed using a standardized protocol and was then divided into aliquots and stored immediately at -80°C. (tumor sample and plasma sample collection) The cervical tumor tissue was obtained from Southern Medical University as an FFPE sample.

### Tumor DNA and CSF cfDNA purification

Cell-free DNA (cfDNA) was isolated from 200–400 µL CSF or plasma using a QIAamp Circulating Nucleic Acid Kit (Qiagen) according to the manufacturer’s instructions. For tumor tissue, DNA was isolated using an AllPrep DNA/RNA Mini Kit according to the manufacturer’s protocol. The isolated DNA concentration was measured using a Qubit 4 Fluorometer with the Qubit dsDNA High Sensitivity Assay Kit (Thermo Fisher Scientific).

### WGBS library preparation

WGBS analysis was used to access the genome-wide DNA methylation profile. The cfDNA WGBS libraries were generated using a Pico Methyl-Seq Library Prep Kit (Zymo Research). Briefly, cfDNA was mixed with 0.1% unmethylated λ-bacteriophage DNA (w/w) (NEB), followed by sodium bisulfite conversion. The bisulfite-converted DNA was then annealed with random primers for initial amplification, followed by adaptor ligation and final amplification with Illumina TruSeq indices. Constructed libraries were run on a 2% agarose gel to assess size distribution, and the library concentration was measured using a Qubit 4 Fluorometer with a Qubit dsDNA High Sensitivity Assay Kit. Normalized libraries were pooled at an equimolar ratio and sequenced on NovaSeq 6000 (Illumina).

### LiBis implementation

LiBis is available on GitHub (https://github.com/Dangertrip/LiBis). LiBis was developed in Python 3.6 and integrated with published software for infrastructure functionalities. In the clipping mode, reads discarded in first-round mapping were clipped using a sliding window according to the user-defined settings for window length and distance from the previous window start base (stride). Clipped read fragments are mapped by BSMAP to find uniquely mapped fragments. Uniquely mapped fragments which are remapped contiguously to the reference genome are concatenated to form rescued fragment candidates. Furthermore, candidates with low read length are discarded to avoid false positives. The minimum read length cutoff which can be adjusted by users is set to 46 base pairs in experiments (**Figure 2B**). For the first-round mapping and remapping, BSMAP used ‘-S 123 -n 1 -r 0 -U’ as setting parameters for uniquely mapped reads and reproducible results. Recombination of contiguously overlapped mapped fragments was implemented according to the following rules to reduce the possibility of false positives: 1) fragments must be strictly continuous, such that the distance between two fragments aligned on the genome must equal the distance between the two sequences on the read; 2) only one mismatch is allowed on each fragment; 3) if, after the recombination, a read overlaps two genomic fragments, the program will select the longest fragment with the fewest mismatches; and 4) all recombined fragments that do not exhibit read overlap are kept. To achieve better time and space complexity, the program records the number of mismatches on the last ‘S’ base pair to accelerate the computation. ‘S’ stands for the length of the stride. The sliding window in the visualization module was achieved through “bedtools makewindows”.

An HTML report template was provided in the pipeline. Datatable and JQuery were imported for data representation. Features of mapping and methylation calling were extracted from the original reports of the corresponding programs to a formatted text file for visualization. Figures were generated during the computation by the Matplotlib package in Python. Principal component analysis plotting used methylation signals in sliding windows along the reference genome.

### Simulation methods

The random nucleotide was generated by the random function in Numpy. The length of the real fragment in the middle was 110 bp from hg38. All cytosines were randomized as unmethylated or methylated. The chance of being methylated was equal to the methylation ratio of the corresponding CpG site in human embryonic stem cells. The random heads of reads had a random length (m) of 1–40 base pairs. The length of the random tail of each read was (40-m) base pairs. The genomic position of the middle part of the read was selected by a two-step randomization that generated the chromosome name and the starting point. For the fully randomized dataset, each nucleotide (A, C, T, or G) had an equal probability to fill each position.

### WGBS data processing

All WGBS data mapped by first-round mapping with BSMAP are described as “without LiBis”, and WGBS data mapped by LiBis (i.e., clipped mapping followed by BSMAP) are described as “with LiBis”. scBS-map was applied with default parameters. LiBis used 40 as the window size, five as the stride, and 45 as the filter length. Picard (http://broadinstitute.github.io/picard) was used to remove PCR duplicates from both first round BSMAP results and second round LiBis rescued reads before analysis in processing clinical samples. The smoothed scatterplots (geneplotter in R package) used the CpG sites in common between the two samples as input. Pearson correlations were calculated using the R cor function. Boxplots were plotted using the Python package seaborn. Fixed length trimming utilized Trim-Galore with ‘--clip_R1’ and ‘--clip_R2’. Single cell whole genome bisulfite sequencing data is at the Gene Expression Omnibus database under accession number GSE56879. Data used to compute correlations between LiBis-specific CpGs and public tumor samples are from the GBM tumor sample methylome at GSE121721(17).

### Virus insertion site identification

To identify virus insertion sites by WGBS data, we considered “fusion reads” as supportive evidence. Fusion read is a read which has two nonoverlapping fragments: one was mapped to a virus genome and the other one was mapped to the human genome. Each virus insertion fragment had two insertion sites which can be reflected by mapped fusion read fragments (Figure 5C). When the human fragment was mapped to the Watson strand, the order of fragments on the genome was identical to the order of fragments on the fusion read. When the human fragment was mapped to the Crick strand, the order was reversed. We used the order of fragments on the genome after adjustment by strand information to decide where the mapped reads belong. The left edge of the insertion site was considered as the rightmost base of human fragments when human fragments were on the front of the mapped reads. The right edge was considered as the leftmost human base of reads having a virus fragment on the front. If and only if multiple fusion reads with the same order existed at the same edge and the distance between the leftmost/rightmost bases on edge was less than 5 base pairs, those reads were selected to be supportive reads for the edge. The supported edge was considered as the rightmost/leftmost base of all human fragments. Reported insertion sites in this paper also follow two requirements: 1. Two or more fusion reads support the same insertion site. 2. Position of all rightmost bases was required to be on the left of all leftmost bases.

## Results

### LiBis: a highly cost-efficient, ultrasensitive alignment method for low-input bisulfite sequencing

LiBis is an integrated Python package for processing low-input WGBS data, such as cfDNA methylome and single-cell DNA methylome sequencing data. FastQC, Trim-galore, BSMAP, mcall, and bedtools are integrated into LiBis for quality control, adaptor trimming, read mapping, methylation calling, and functional analysis respectively (13,18-20). The LiBis toolkit contains three modules: a preprocess module for quality control and adaptor trimming, a compute module for dynamic clipping and mapping, and a visualization module for report generation (**Figure 1A, Figure S1**).

**Figure 1.**
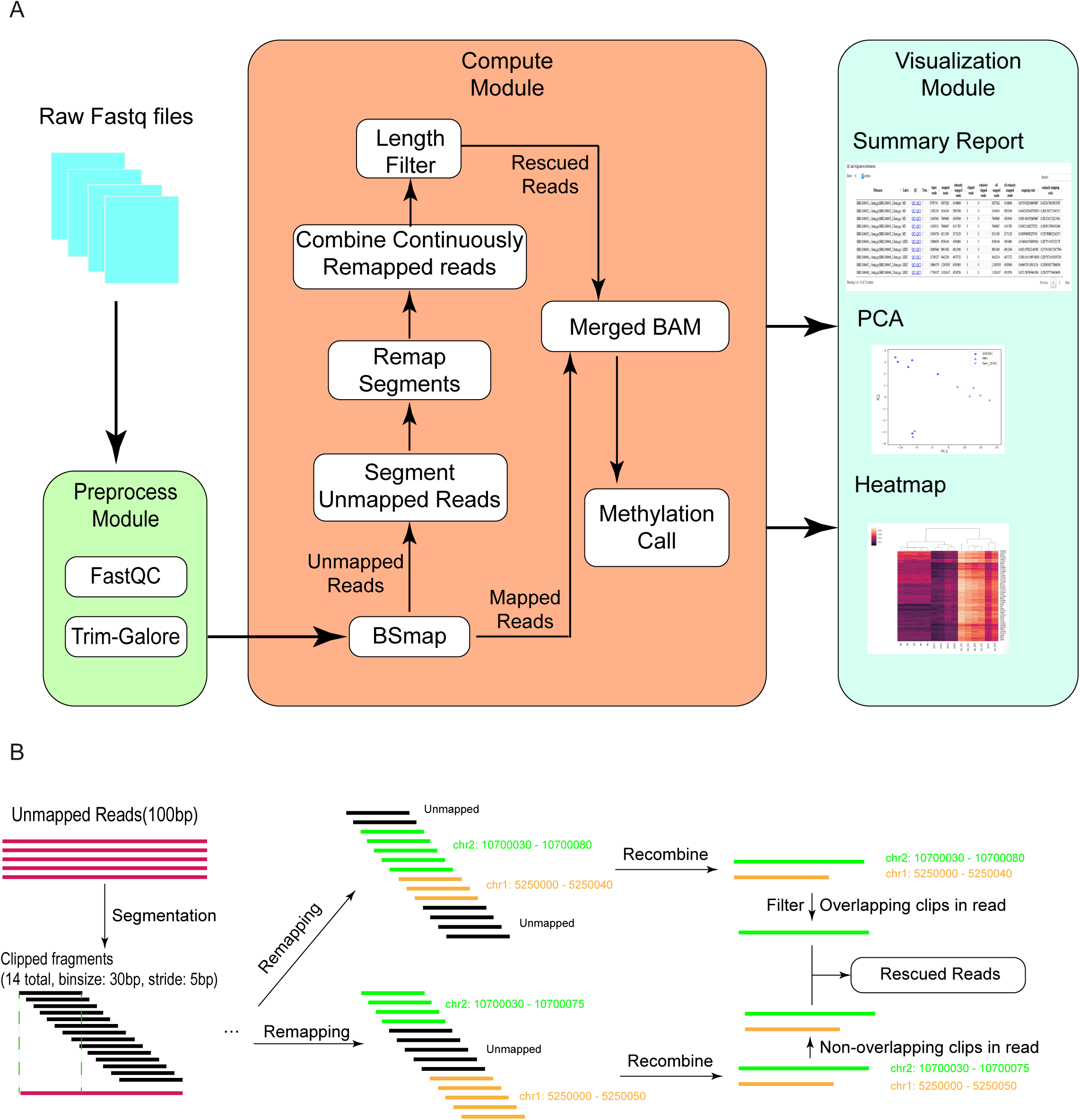
Development of LiBis (low-input bisulfite sequencing alignment). A. LiBis overview. B. Details of the LiBis rescue procedure, including clipping initially unmapped reads, remapping clipped fragments, and recombining contiguous fragments.

**Figure 2.**
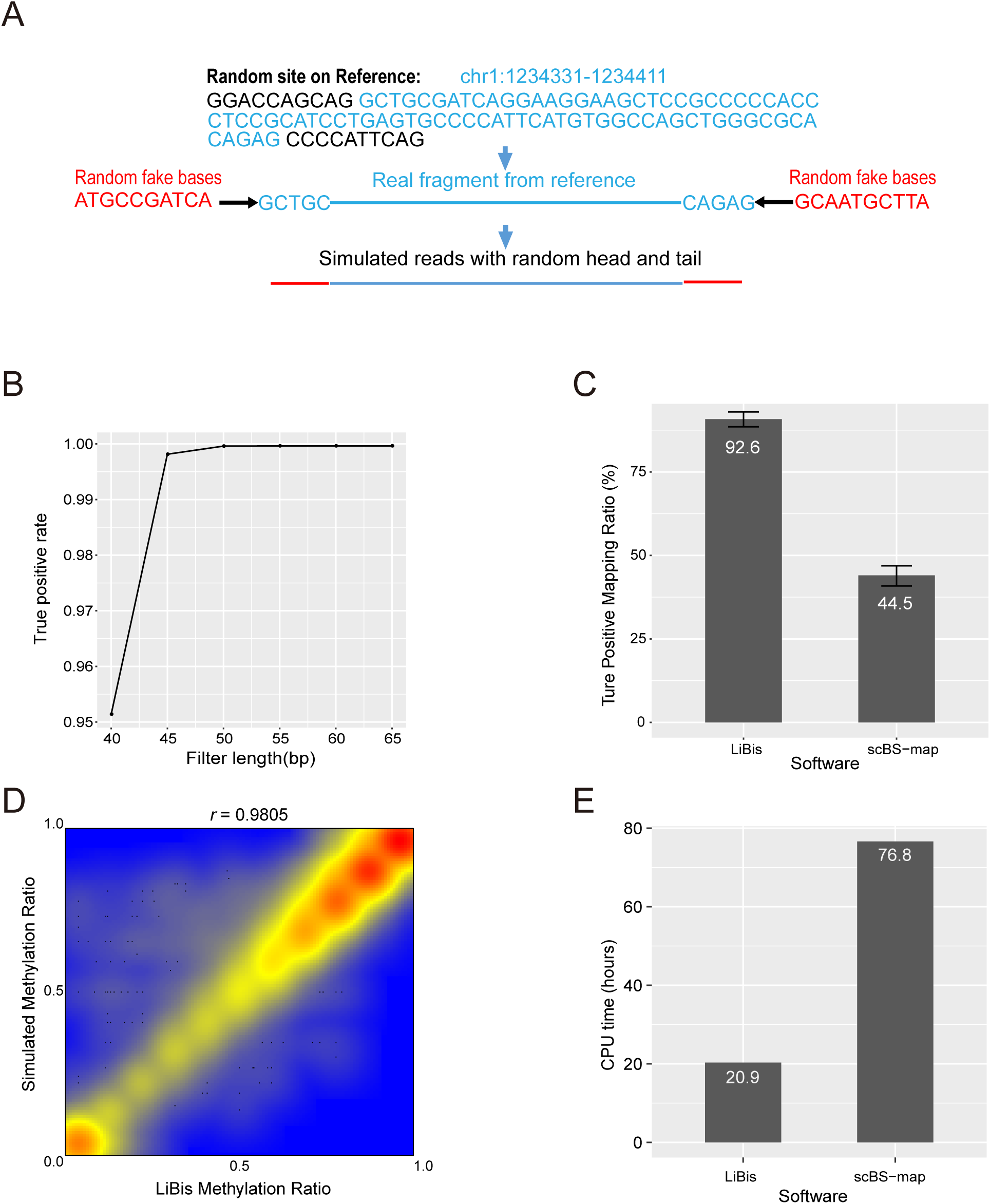
LiBis shows a high discovery rate, high sensitivity, high accuracy, and fast efficiency *in silico*. A. Scheme for generating simulated reads with random heads and tails. B. True positive rates of LiBis using different filter lengths. C. Mapping ratios of simulated reads by LiBis or scBS-map. D. Methylation ratio correlation between simulation ground truth and LiBis. E. CPU times for processing 10 million reads by LiBis or scBS-map.

To improve the mapping efficiency for low-input bisulfite sequencing data, we applied a clipping strategy within the compute module to eliminate random base contamination, as follows. First, LiBis maps the trimmed raw data with BSMAP and generated a new fastq file containing unmapped reads. Second, LiBis clips all unmapped reads using a sliding window of a specific width and step size as defined by the user. Third, LiBis remaps all clipped read fragments and keeps only uniquely mapped fragments for subsequent recombination. During recombination, fragments derived from the same unmapped read are recombined only if they are remapped contiguously to the reference genome, with the distance between two adjacent fragments being the step size. Recombined fragments are required to be above a minimum length to reduce the likelihood of a false-positive alignment. For reads with multiple candidate recombined clipped fragments that overlap, such as that illustrated in the top half of **Figure 1B**, the recombined clip with the greatest mapping confidence (i.e., the longest clip with least mismatches) is kept as a rescued read. If the recombined clipped fragments do not overlap each other, such as the two clips in the bottom half of **Figure 1B**, all recombined clipped fragments will be kept. Through the remapping and recombination steps mentioned above, reads discarded in first-round mapping can be rescued.

The LiBis workflow for bisulfite sequencing data involves five steps. In step 1, the raw reads are examined for quality control by FastQC. FastQC allows us to assess quality features including base quality, base content, and duplication level. These features reveal the overall quality of the library, amplification, and sequencing. Results of FastQC are aggregated in the final report. In step 2, the reads are trimmed by trim-galore, which removes sequencing adaptors and low-quality reads. If random priming was used to generate the library, trimming is recommended but will not remove random priming-associated amplification artifacts. Step 3 is read mapping by the compute module, which combines initial mapping with the dynamic clipping and remapping strategy to improve the cost efficiency. In step 4, the methylation ratio of CpGs is called by mcall from the MOABS program. In step 5, the data is visualized as various figures. After LiBis analysis, an overview webpage presents summaries of all input samples, heatmaps, and principal component analysis results (**Figure 1A**).

LiBis requires two types of input files, namely the reference genome sequence file in fasta format and the fastq files containing raw reads. For small numbers of samples, users can use command line to run LiBis. Additionally, we developed a config file reader for large numbers of samples. The LiBis output contains bam files, methylation ratio files, quality control results, stats files and an integrated HTML report.

### LiBis improves mapping ratios and mapping sensitivity in simulated data

Three simulation datasets were generated randomly and independently *in silico* with 10 million 150 base pair length reads for each dataset. Each 150 bp length read is composed of a 110 bp DNA sequence (randomly cut from hg38 human genome) and random sequences on both ends of the 110 bp genome DNA sequences. The random heads and tails simulated contamination introduced by the random priming process, which cannot be fully removed by traditional trimming methods such as trim-galore (**Figure 2A**). LiBis’ mapping ratio is associated with the minimum length of rescued reads. We investigated the relationship between true positive rates and the filter length. As shown in Figure 2B, the true positive rate was saturated when 45 base pairs was set as the minimum rescued length filter. All simulated datasets were mapped using LiBis and scBS-map as the state of the art (16). LiBis achieved a mapping ratio (92.6%) which is higher than scBS-map (44.5%) (**Figure 2C**). Regarding sensitivity, an event was defined as true positive if the interval of the simulated read on the genome included its mapping interval. For the mapped reads, LiBis achieved over 99.9% sensitivity on average, which was significantly higher than the 92.9% sensitivity of scBS-map (**Figure S3**). The DNA methylation ratios of CpGs identified by LiBis also showed a high correlation (r=0.98) with the ground truth for the simulation (**Figure 2D**), indicating the high accuracy of LiBis. These results revealed that LiBis achieves high detectability and high sensitivity in simulation datasets.

Regarding efficiency, our results showed that LiBis used only 20.9 CPU hours to process the simulated dataset (10 million 150 bp length reads), which is comparable to BSMAP (10.6 CPU hours) considering the rescue of contaminated reads from simulation dataset. Compared to other software, LiBis is nearly four times faster than scBS-map (**Figure 2E**).

### LiBis improved mapping ratios and mapping sensitivities in single-cell DNA methylome dataset

To test the efficiency of LiBis using real data, we applied LiBis on single cell mouse ESC WGBS samples from a public dataset (GSE56879). We observed that the mapping ratio for the scWGBS samples increased by as much as 24% with LiBis compared to BSMAP (**Figure 3A**). A large number of CpGs were specifically discovered by LiBis, but failed to be detected by BSMAP (**Figure 3B**). To further interpret these improvements regarding its effects on downstream analysis, we defined CpGs that were covered at least 5 times as informative CpGs. LiBis recovered as many as 70% more informative CpGs compared to BSMAP, dramatically increasing the number and depth of usable CpG sites (**Figure 3C**). For the CpG sites specifically discovered by LiBis, our result shows that these CpGs had relatively higher methylation ratios compared to the initially discovered CpGs (**Figure S3A**), indicating that reads from highly methylated regions may suffer more severe contamination from random priming, which may partially explain the lower methylation ratios detected in libraries built using random priming versus in libraries prepared using pre-bisulfite sequencing methods (10).

**Figure 3.**
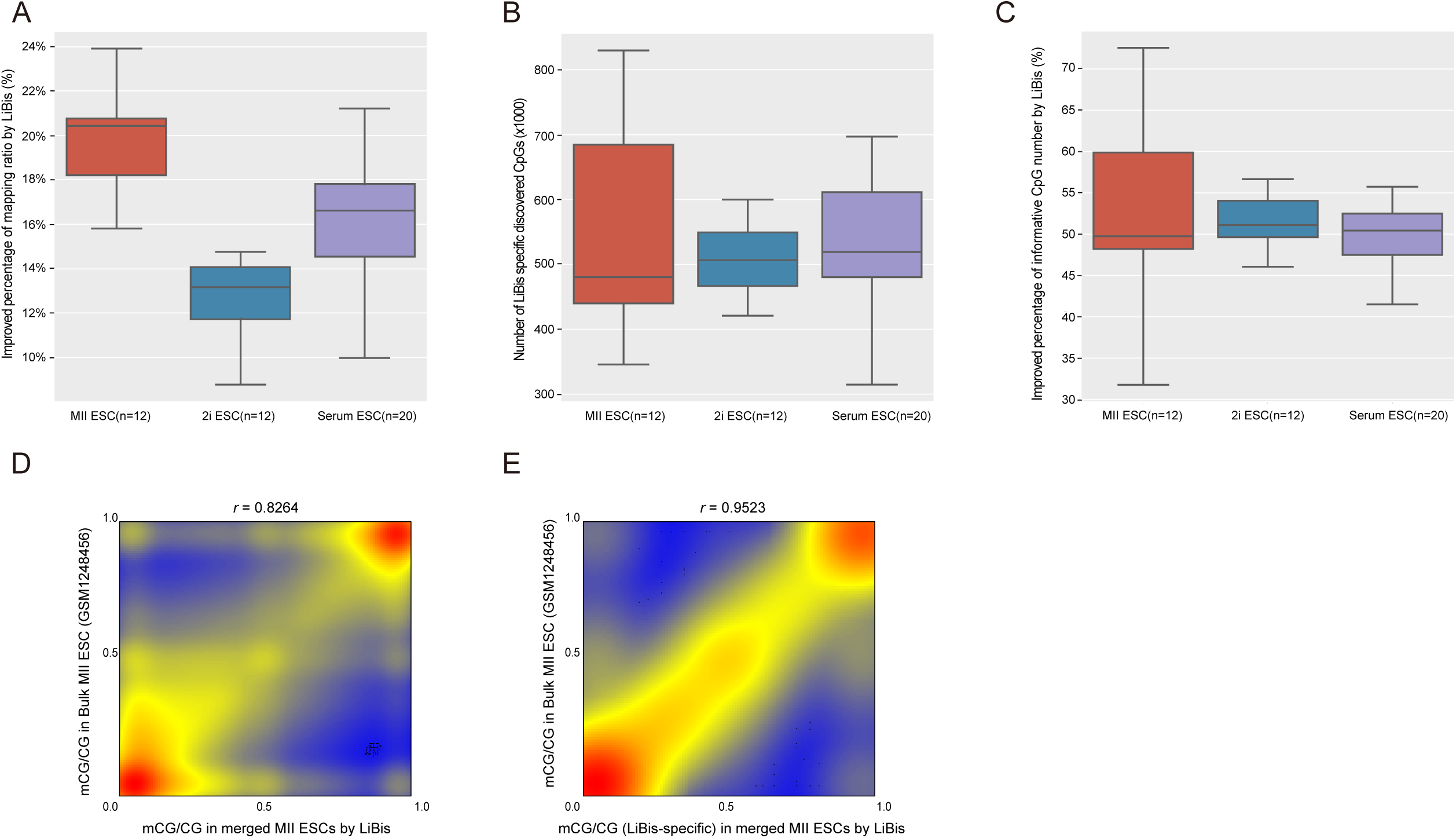
LiBis improves both the efficiency and the precision of methylation measurements in scWGBS samples. A–C. Percentage change in the mapping ratio, the number of CpGs recovered only by LiBis-rescued reads compared to BSMAP, and the number of informative CpGs (depth ≥5) D. Correlation of the methylation ratio (averaged for 1-kb windows) between bulk and merged single cell methylomes. E. Methylation ratio correlation between bulk and merged methylomes at regions specifically discovered by LiBis.

To estimate the accuracy of the DNA methylation ratios recovered by LiBis, we performed a comparison between DNA methylation ratios generated by LiBis using mixed scWGBS data (LiBis-MII ESCs-scWGBS) and the DNA methylation ratios generated by BSMAP using bulk methylome data (BSMAP-MII ESCs-bulk) from the same cell resources. We observed a high correlation in the DNA methylation ratio (Pearson correlation r = 0.83) between LiBis-MII ESC-scWGBS and BSMAP-MII ESCs-bulk on the 2.5 million common detected CpGs. (**Figure 3D, S3B**). This result indicated that LiBis can identify CpGs with accurate DNA methylation ratios. There are 187,804 out of 2.5 million CpGs that were specifically recovered by LiBis in merged single cell samples, which also highly correlated with BSMAP-MII ESCs-bulk (Pearson correlation r=0.95) (**Figure 3E**). Additionally, the methylation ratio difference between the BSMAP bulk methylome and the merged scWGBS methylome by BSMAP or LiBis was subtle and comparable with BSMAP’s results (**Figure S3C**). These results strongly demonstrated that the CpG DNA methylation ratios from LiBis rescued reads are accurate and beneficial for downstream analysis.

### LiBis improved the data efficiency of tumor and cfDNA WGBS experiments with random priming

To verify the capability of LiBis for rescuing bisulfite sequencing data from low input WGBS (such as cfDNA samples), we performed WGBS on six clinical samples and processed the sequencing data using LiBis: 1). Two cerebrospinal fluid (CSF) cfDNA samples from glioblastoma patients; 2). Two plasma cfDNA samples from glioblastoma patients; 3). Two genomic DNA samples from glioblastoma tumor tissues. We adopted random priming-based WGBS to generate libraries of all collected samples (9). To validate the functionality of LiBis, we first compared the traditional fixed length trimming method and LiBis (**Figure 4A**). Although the fixed length trimming method can improve the mapping ratio, there was only a slight change in the base pair usage rate (right Y-axis). LiBis can significantly improve both the mapping ratio and base pair usage over the traditional fixed length trimming method. Furthermore, by analyzing the base content of the sequences, we found that our samples prepared using the random priming library method showed a strong C-bias at the beginning of the reads. This observation was also reported in papers using single-cell bisulfite sequencing due to the dNTP in random priming (11,21). LiBis can effectively eliminate the C-bias by dynamically removing random artifactual bases in both ends of the read (**Figure 4B**).

**Figure 4.**
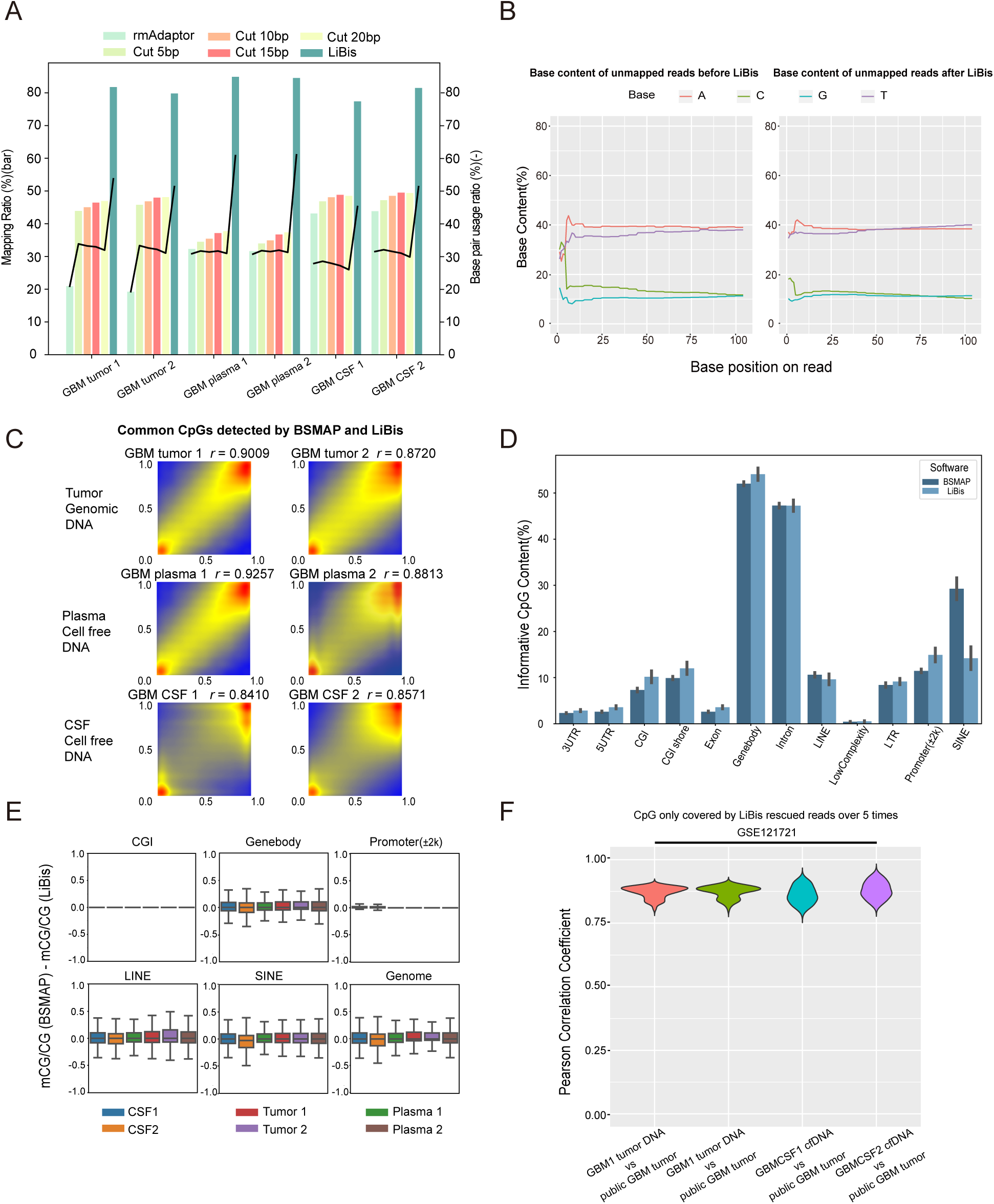
LiBis improves the cost efficiency of clinical tumor and cfDNA WGBS data using the random priming method. A. Improvement comparison between LiBis and fixed length trimming. B. Base content distribution by BSMAP and LiBis. Content bias can be eliminated by LiBis. C. Correlation of the methylation ratios between the BSMAP mapped reads and the rescued reads in LiBis. The correlation coefficients are listed on the top of the figures. D. Distribution of informative CpGs as determined by BSMAP and LiBis. E. Minor methylation differences between BSMAP and LiBis were found in common CpGs in genomic regions and the global genome. F. Correlations of methylation ratios between LiBis-specific CpGs and collected patient methylomes from public database.

**Figure 5.**
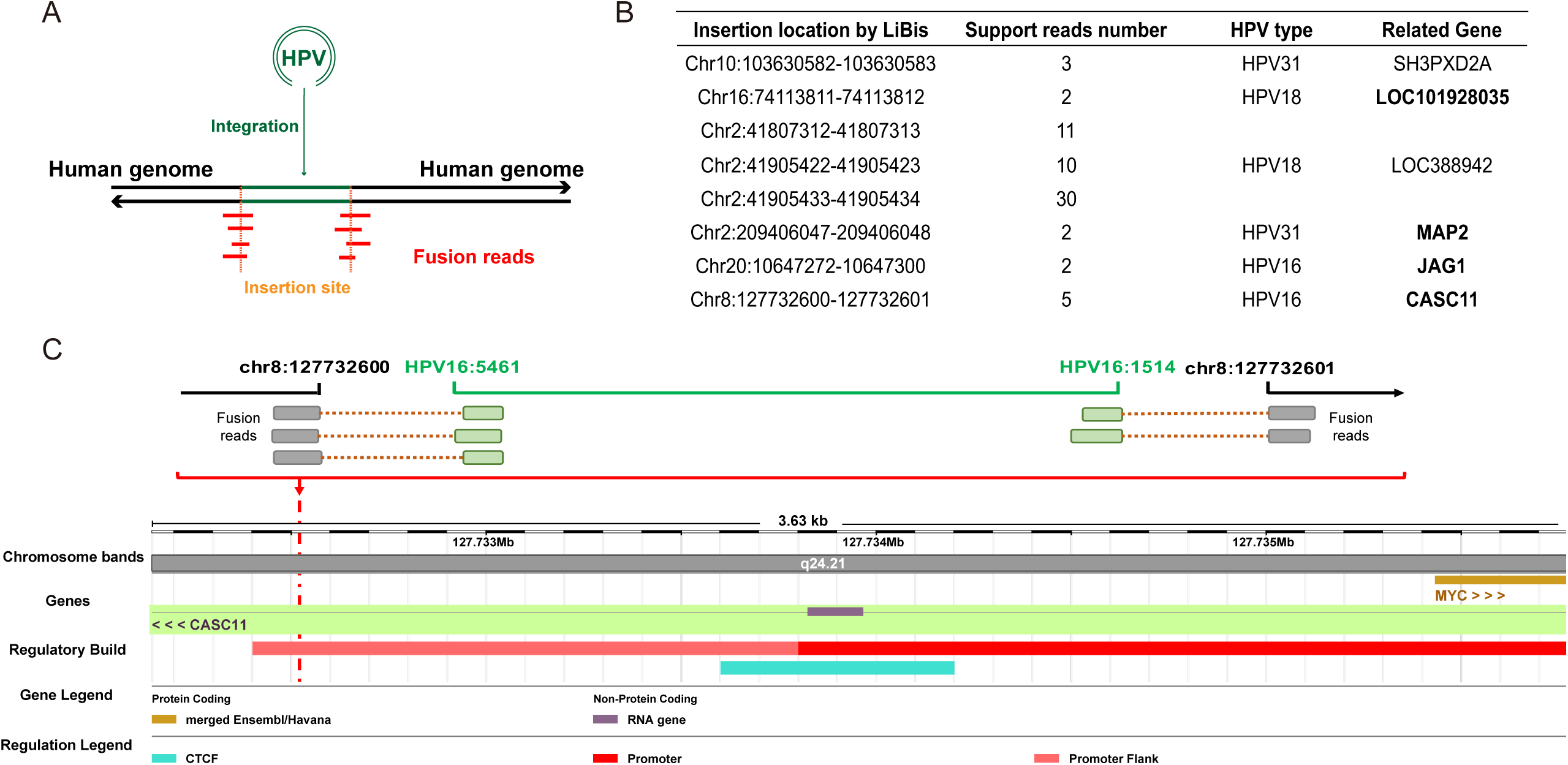
LiBis identifies HPV insertion sites in cervical cancer WGBS experiments using a traditional pre-bisulfite library method. A. Scheme of HPV insertion. B. HPV insertion sites in cervical cancer identified by LiBis. C. Visualization of an identified HPV insertion site.

Next, to measure the similarity in the DNA methylation ratios of each CpG comparing reads recovered by LiBis and using the end-to-end mapping reads, we performed correlation analysis. We observed a high correlation coefficient that ranged from 0.84 to 0.92 (average 0.88) between the DNA methylation ratios from the end-to-end mapping and the LiBis clipped mapping on the CpGs covered least 10 times by both methods (**Figure 4C)**. LiBis recovered informative CpGs share a similar distribution in different genomic regions while having a relatively low recovery rate in repeat regions (**Figure 4D**). The reason for the low number of LiBis rescued reads in repeat regions is that short fragments may lead to multiple mapping on repeat regions, which leads to a lower unique mapping ratio on repeat regions. In addition, the majority of the CpGs detected in common between BSMAP and LiBis have a DNA methylation ratio difference less than 0.2 across genome elements. (**Figure 4E**). CpGs at low methylated regions such as CpG islands or promoters showed no methylation ratio difference because as the methylation ratio distribution becomes more skew, the confidence interval narrows. These results further showed that LiBis faithfully recovered CpG DNA methylation ratios. CpGs uniquely recovered by LiBis had DNA methylation ratios that were highly correlated with those from public tumor tissue methylome databases (average Pearson coefficient r=0.87) (**Figure 4F**). These results strongly demonstrated that LiBis can efficiently and correctly extract maximum DNA methylation information from low-input bisulfite sequencing data in clinical samples.

### LiBis can identified human papillomavirus integration sites in cancer using WGBS data

To further explore the capability of LiBis to discover genomic patterns, we collected one cervical cancer tumor sample and prepared a WGBS library using a traditional protocol to prepare the sequencing library. The mapping ratio improved from 84% with BSMAP to 90% with LiBis using the 150 bp paired-end sequencing strategy. Although the improvement was relatively small compared to that observed for the cfDNA WGBS data, we confirmed that WGBS together with LiBis could identify DNA fusion sites, in particular the human papillomavirus (HPV) DNA-human DNA fusion sites in this case study. In cervical cancer, HPV DNA can cleave human DNA and insert itself into the human genome in a sense or antisense direction. Such fusion reads containing both HPV and human DNA fragments were identified by LiBis, which identified the insertion spots of HPV (**Figure 5A**). We identified a list of potential insertion sites in this patient sample using LiBis (**Figure 5B**). For each insertion site, we identified the closest gene to find the potential influence of the viral insertion on gene expression. Interestingly, four of the six insertion-proximal genes we identified were previously reported as HPV insertion sites or were differentially expressed in cervical cancer (22-24). For example, one insertion site was located in the intron of the *CASC11* gene and in the flanking promoter region of the *MYC* gene (**Figure 5C**). *CASC11*, a gene that is upregulated in cervical cancer tissue, might activate the WNT/β-catenin signaling pathway to promote cervical cancer, and the *MYC* gene is well known for contributing to tumorigenesis (25). Interestingly, at this insertion site HPV integrates in a non-linear fashion. In addition to identifying HPV integration sites, LiBis enabled the identification of differentially methylated CpGs in both human and HPV genomes using the same pipeline. These results suggest that LiBis could be used to identify the viral integration sites using WGBS data, which will be beneficial for the DNA methylation analysis of viral integration in cancer.

## Discussion

DNA fragments sequenced by post-bisulfite sequencing are reported to have self-priming and chimeric reads, which leads to low efficiency of utilizing sequencing raw data (11,12). In this study, we developed a pipeline called LiBis to reduce this problem in post-bisulfite sequencing by fragmentizing, remapping and recombining unmapped reads. To the best of our knowledge, LiBis is the first integrated pipeline for the processing of low-input bisulfite sequencing data that includes trimming, initial mapping, clipped mapping, methylation calling, and visualization. By dynamically clipping unmapped reads to remove random priming contamination, LiBis is capable of improving the cost efficiency of experiments and the measurement precision. We confirmed that LiBis improved performance in four case studies, namely simulated data, scWGBS data, cfDNA WGBS data from random priming libraries, and tumor WGBS data from random priming libraries. The performance improvement may due to the following reasons. First, random priming, adapter dimers, small fragments, or bisulfite conversion may generate artifactual bases at both ends of the reads. Our approach reduces the proportion of reads culled due to these issues. Second, through dynamic clipping, LiBis could recognize DNA fusions present in the original cell, including gene fusions, insertions of HPV DNA, fusions that arose during the library preparation, and even circular DNA formed by self-priming or other mechanisms.

To maximize the mapping efficiency of post-bisulfite sequencing material, our method shares similar concepts with previous reports. scBS-map applies a local alignment after an end-to-end mapping step to rescue discarded reads (16). Taurus-MH splits the unmapped reads in end-to-end mapping with a fixed pattern to estimate the mappable parts (26). Although all of the current methods use read splitting or soft clipping to rescue unmapped reads, however, our method have several distinct characteristics. All parameters in the clipping process in LiBis such as window length and stride can be self-defined before initiating the mapping process. The simplicity of strategy and parameters in LiBis allows users to adjust the parameters to balance the running time and the rescuing efficiency. For example, users can have a smaller stride and window length when the sequencing quality is unsatisfactory to improve the cost efficiency. Compared to previous methods, LiBis has 2 advantages: 1. LiBis had significantly higher efficiency than the traditional trimming method or other mapping software in processing WGBS data from a random priming library. 2. LiBis is an integrated pipeline including quality control, trimming, mapping, methylation calling and visualization modules, which can achieve one-command processing for WGBS raw data. Finally, all methods including LiBis can be further enhanced by applying parallel computing for mapping processes to query more computational capacity. A more precise iteration of parameters for LiBis can also improve the efficiency batch to batch.

## Conclusions

We developed LiBis, a novel dynamic clipping and mapping method for WGBS of low-input DNA. Using fastq files as input, LiBis outputs a well-organized HTML report of results from the different modules. LiBis significantly improved the cost efficiency for analyzing low-input bisulfite sequencing data by providing a larger number of accurate informative CpGs and increasing the sequencing depth of all CpGs for downstream analysis. LiBis provides accurate methylome information through conservative mapping strategies, as illustrated by the simulation data and the data from the scWGBS and bulk WGBS experiments. Moreover, LiBis can identify DNA fusion features, such as virus insertion sites, from WGBS data.

## Data Access

All raw and processed sequencing data generated in this study are available at the Gene Expression Omnibus database under accession number GSE142241.

## Software Requirements

Project name: LiBis

Operating system(s): Linux or system based on Docker environment

Programming language: Python 3.6

Other requirements: Docker (Optional)

License: GNU

## List of Abbreviations

LiBis: low-input <u>bis</u>ulfite sequencing alignment
cfDNA: cell-free DNA
WGBS: whole-genome bisulfite sequencing
CpG: cytosine nucleotide followed by a guanine nucleotide
ctDNA: circulating tumor DNA
HTML: HyperText Markup Language
HPV: Human papillomavirus

## Acknowledgements

We are grateful to the Texas A&M University Brazos HPC cluster (brazos.tamu.edu) and the Texas A&M Institute for Genome Sciences and Society (TIGSS) HPC cluster (tigss.tamu.edu) that contributed to the research reported here.

## Author Contributions

DS directed and oversaw the project. YY developed and tested the pipeline and performed the computational experiments. YY and JL performed comprehensive bioinformatics analysis, including data quality control, publicly available data collection, and integration analysis. ML and YH optimized CSF ctDNA sequencing library preparation and performed high-throughput sequencing. SZ, XNL, and LG collected the samples. Jin, Jianfang, and Mutian provided intellectual input. YY and DS wrote the manuscript. All authors contributed to discussion, program testing, and the writing of the manuscript.

## Supplementary Figure Legends

**Figure S1**. Example of the final report generated by the LiBis visualization module. Top panel, table of basic statistics and HTML links to FastQC figures. Bottom panel, principal component analysis scatter plot and heatmap.

**Figure S2**. True positive rates of mapped reads by LiBis or scBS-map.

**Figure S3**. A. Regions specifically discovered by LiBis have a higher methylation ratio than other regions. B. Merged methylomes generated with LiBis or without LiBis exhibit nearly identical methylation ratio distributions over gene regions. All genes in the genome were stretched to 1 kb and aligned to the interval from Start to End on the X axis. The Y axis is the methylation ratio. The bulk methylome by BSMAP is in red, the merged methylome with or without LiBis is in green or blue, respectively. C. The distributions of methylation ratio difference between bulk and merged methylomes are similar in BSMAP and LiBis.

