## Supplemental Figure 1 for "LiBis: An ultrasensitive alignment method for low-input bisulfite sequencing"

QC and Alignment information:

Show 10 entries

Search:

| Filename | Label | QC | Trim | Input reads | mapped reads | uniquely mapped reads | clipped reads | uniquely clipped reads | all mapped reads | all uniquely mapped reads | mapping ratio | uniquely mapping ratio |
|---|---|---|---|---|---|---|---|---|---|---|---|---|
| SRR1248451_1.fastq.gz,SRR1248451_2.fastq.gz | MII | QC1,QC2 | / | 9797740 | 6597262 | 4146868 | 0 | 0 | 6597262 | 4146868 | 0.6733452816669967 | 0.4232474019518787 |
| SRR1248452_1.fastq.gz,SRR1248452_2.fastq.gz | MII | QC1,QC2 | / | 13921551 | 6184434 | 5083548 | 0 | 0 | 6184434 | 5083548 | 0.44423455403783674 | 0.3651567271491517 |
| SRR1248453_1.fastq.gz,SRR1248453_2.fastq.gz | MII | QC1,QC2 | / | 15447641 | 7849660 | 4345949 | 0 | 0 | 7849660 | 4345949 | 0.5081461952669667 | 0.2813341532211941 |
| SRR1248454_1.fastq.gz,SRR1248454_2.fastq.gz | MII | QC1,QC2 | / | 14038511 | 7696067 | 4191783 | 0 | 0 | 7696067 | 4191783 | 0.548211060275552 | 0.2985917096193464 |
| SRR1248455_1.fastq.gz,SRR1248455_2.fastq.gz | MII | QC1,QC2 | / | 15544734 | 6311109 | 5173129 | 0 | 0 | 6311109 | 5173129 | 0.405996590227919 | 0.3327898052163517 |
| SRR1248464_1.fastq.gz,SRR1248464_2.fastq.gz | 2iESC | QC1,QC2 | / | 15966034 | 8536146 | 4594884 | 0 | 0 | 8536146 | 4594884 | 0.5346441076099424 | 0.2877911947325178 |
| SRR1248465_1.fastq.gz,SRR1248465_2.fastq.gz | 2iESC | QC1,QC2 | / | 18065844 | 8891060 | 4912346 | 0 | 0 | 8891060 | 4912346 | 0.4921475022146765 | 0.27191345170477504 |
| SRR1248466_1.fastq.gz,SRR1248466_2.fastq.gz | 2iESC | QC1,QC2 | / | 21749327 | 8442254 | 4475732 | 0 | 0 | 8442254 | 4475732 | 0.38816161989748005 | 0.20578714918397245 |
| SRR1248467_1.fastq.gz,SRR1248467_2.fastq.gz | 2iESC | QC1,QC2 | / | 19066379 | 12367855 | 4385960 | 0 | 0 | 12367855 | 4385960 | 0.6486735105811124 | 0.2300363377860054 |
| SRR1248468_1.fastq.gz,SRR1248468_2.fastq.gz | 2iESC | QC1,QC2 | / | 17740157 | 11924167 | 4518556 | 0 | 0 | 11924167 | 4518556 | 0.6721567909461004 | 0.2547077796436638 |

Showing 1 to 10 of 15 entries

Previous 1 2 Next

Cluster figures:

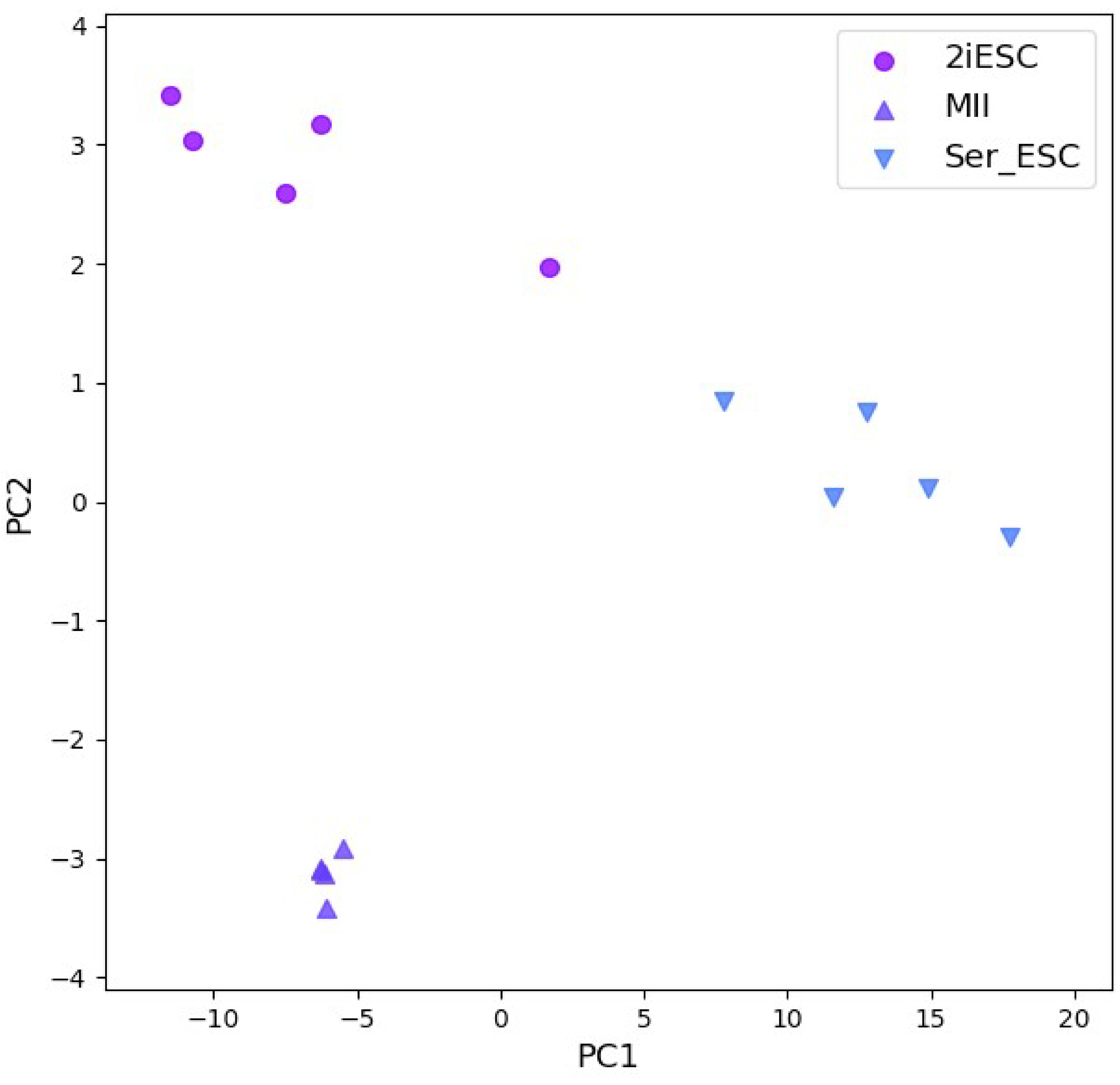


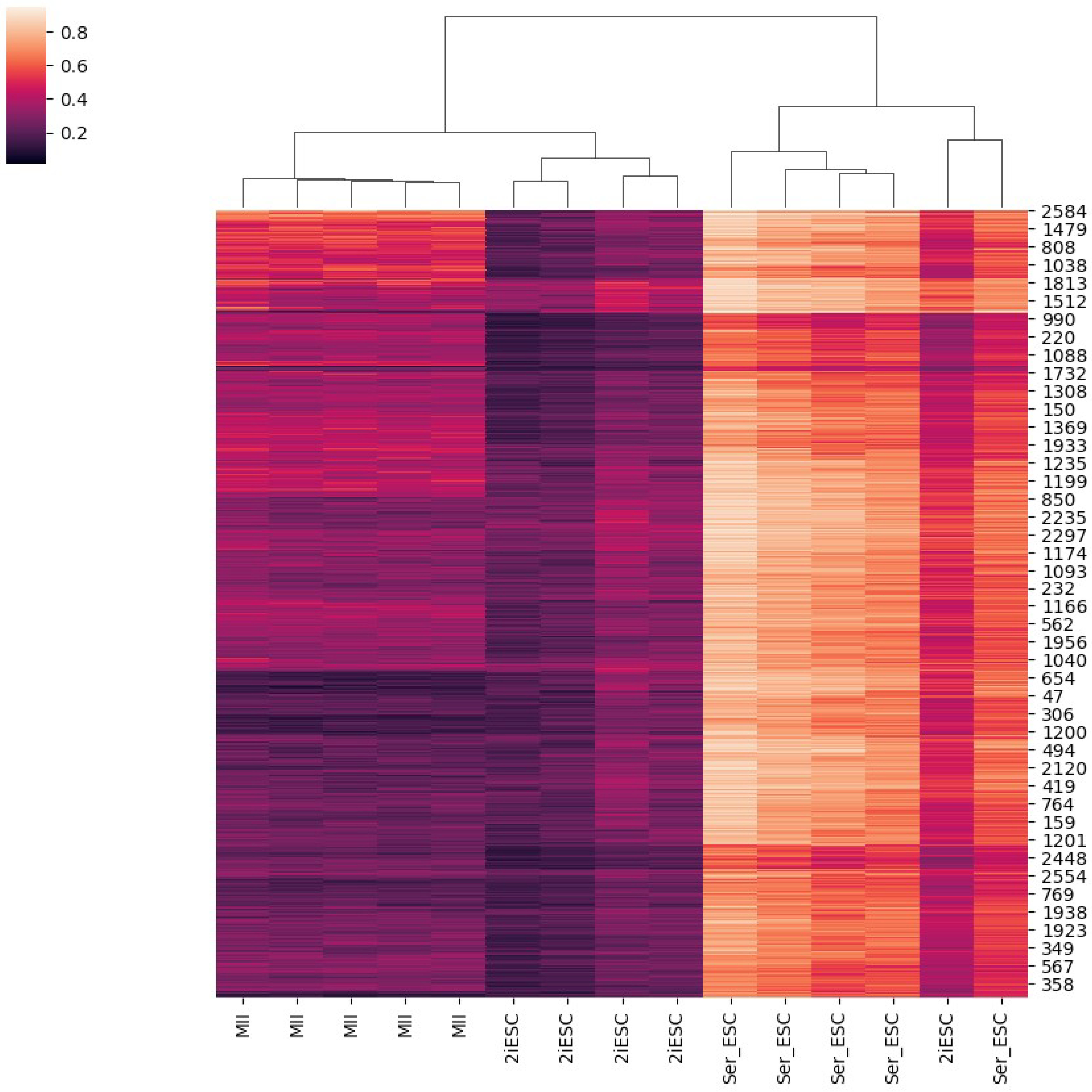
