## Supplementary figures and images for "LiBis: An ultrasensitive alignment method for low-input bisulfite sequencing"

### Supplemental Figure 2

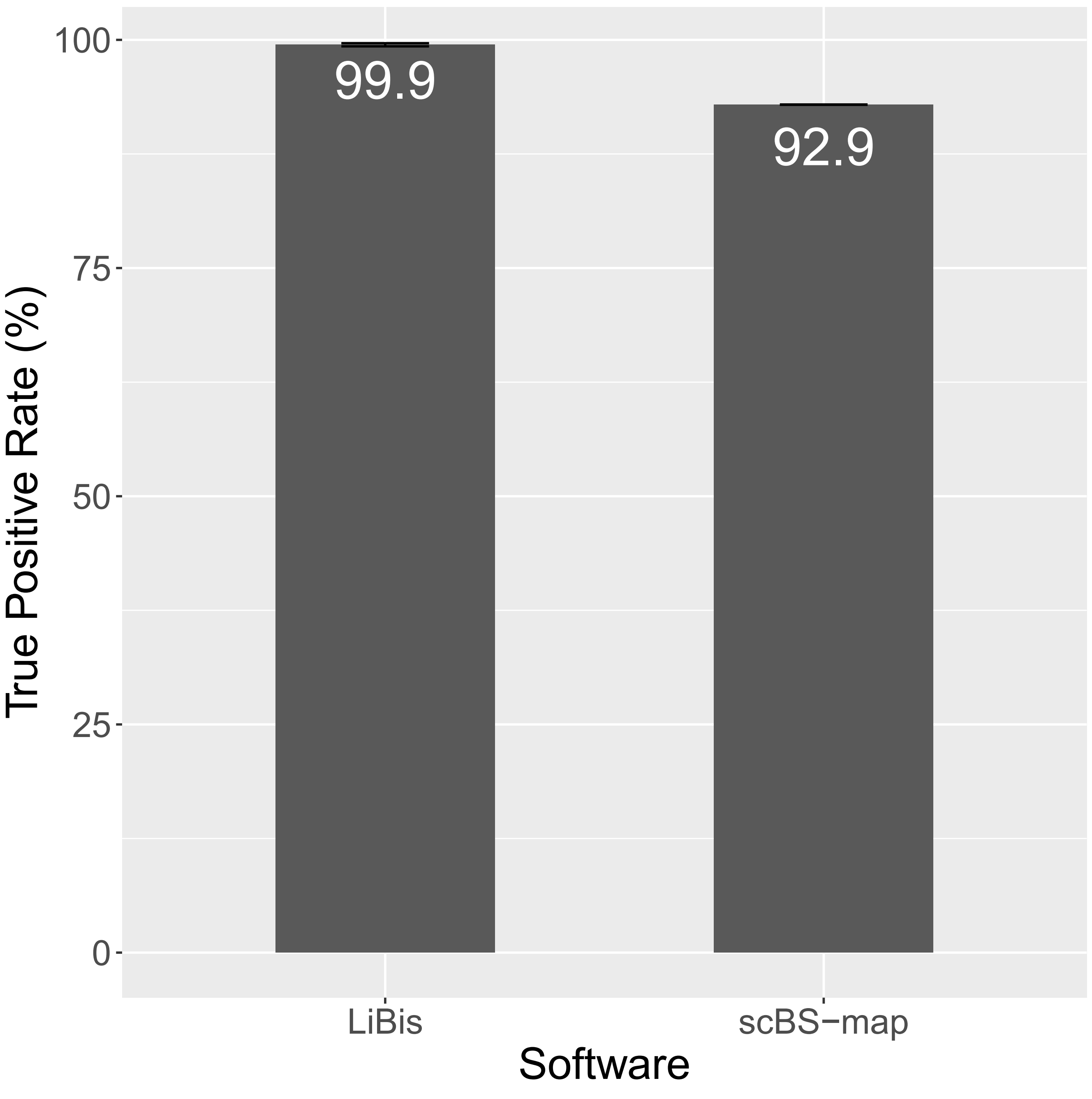

### Supplemental Figure 3

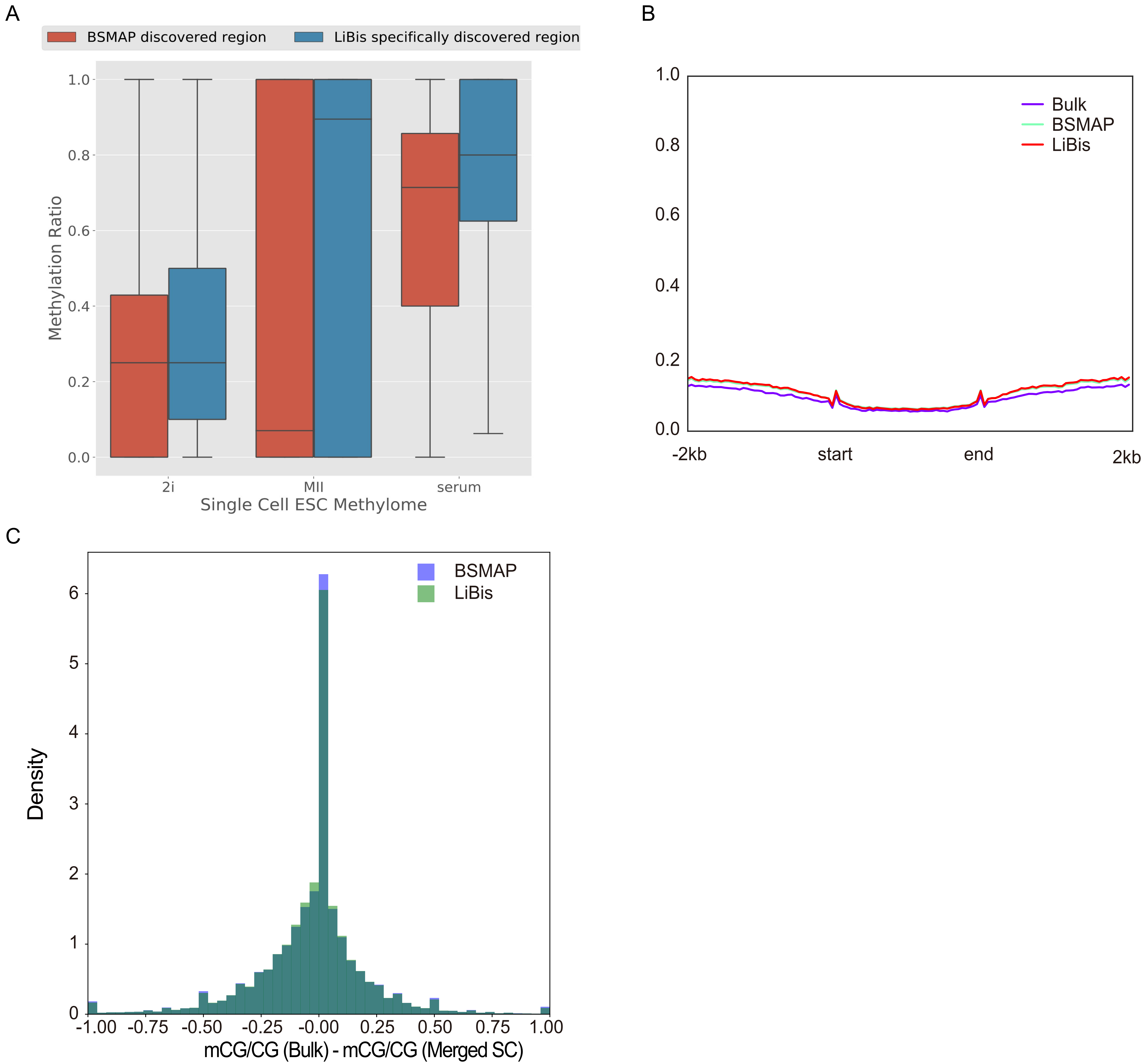

### Supplemental Figure 4

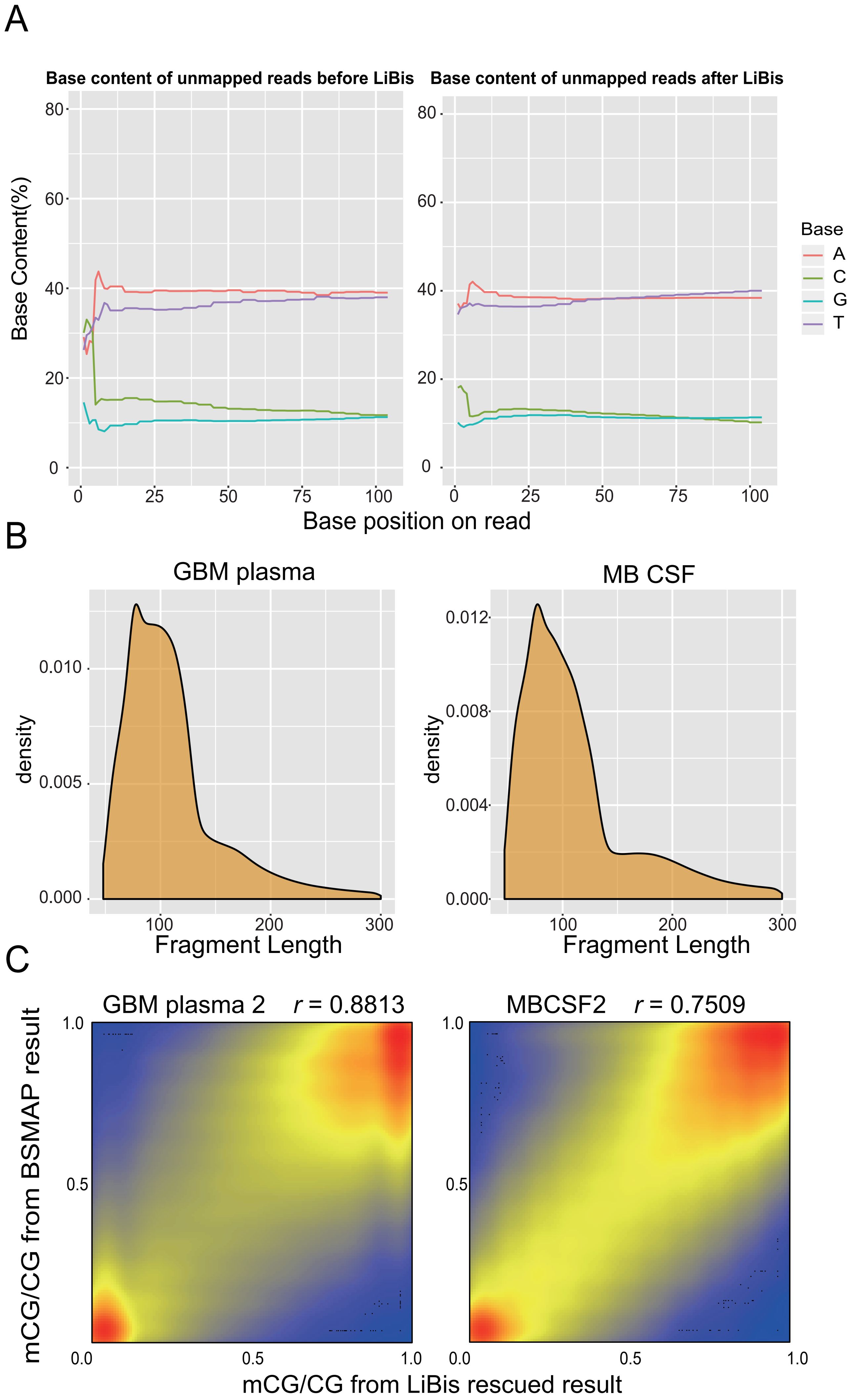
